# Robustness of working memory to prefrontal cortex microstimulation

**DOI:** 10.1101/2025.01.14.632986

**Authors:** Joana Soldado-Magraner, Yuki Minai, Byron M. Yu, Matthew A. Smith

**Affiliations:** Department of Electrical and Computer Engineering, Carnegie Mellon University, Pittsburgh 15213, Pennsylvania, USA; Department of Biomedical Engineering, Carnegie Mellon University, Pittsburgh 15213, Pennsylvania, USA; Machine Learning Department, Carnegie Mellon University, Pittsburgh 15213, Pennsylvania, USA.; Center for the Neural Basis of Cognition, Carnegie Mellon University and University of Pittsburgh, Pittsburgh 15213, Pennsylvania, USA; Neuroscience Institute, Carnegie Mellon University, Pittsburgh 15213, Pennsylvania, USA

**Author notes:** Correspondence: Joana Soldado-Magraner. equal contribution (Byron M. Yu, Matthew A. Smith).

**Keywords:** Dorso-lateral prefrontal cortex, neural perturbations, population activity, dimensionality reduction

## Abstract

Delay period activity in the dorso-lateral prefrontal cortex (dlPFC) has been linked to the maintenance and control of sensory information in working memory. The stability of working memory related signals found in such delay period activity is believed to support robust memory-guided behavior during sensory perturbations, such as distractors. Here, we directly probed dlPFC’s delay period activity with a diverse set of activity perturbations, and measured their consequences on neural activity and behavior. We applied patterned microstimulation to the dlPFC of monkeys implanted with multi-electrode arrays by electrically stimulating different electrodes in the array while the monkeys performed a memory-guided saccade task. We found that the microstimulation perturbations affected spatial working memory-related signals in individual dlPFC neurons. However, task performance remained largely unaffected. These apparently contradictory observations could be understood by examining different dimensions of the dlPFC population activity. In dimensions where working memory related signals naturally evolved over time, microstimulation impacted neural activity. In contrast, in dimensions containing working memory related signals that were stable over time, microstimulation minimally impacted neural activity. This dissociation explained how working memory-related information could be stably maintained in dlPFC despite the activity changes induced by microstimulation. Thus, working memory processes are robust to a variety of activity perturbations in the dlPFC.

**Significance statement:** Memory-guided behavior is remarkably robust to sensory perturbations such as distractions. The dorso-lateral prefrontal cortex (dlPFC) is believed to underlie this robustness, given that it stably maintains working memory-related information in the presence of distractors. Here, we sought to understand the extent to which dlPFC circuits can robustly maintain working memory information during memory-guided behavior. We found that behavior was robust to microstimulation perturbations in dlPFC, and that working memory signals were stably maintained in dlPFC despite widespread changes in the neural activity caused by the perturbations. Our findings indicate that working memory is robust to direct activity perturbations in the dlPFC, an ability that may be due to the processes that mediate similar robustness in the face of distraction.

## Introduction

When animals are engaged in delayed response tasks that require the maintenance and control of sensory-related information to guide future actions, the brain must store this information in working memory and protect it from interference from other external and internal signals (Katsuki and Constantinidis, 2012; Lorenc et al., 2021; Wang, 2021). For example, in tasks that require remembering the location of a visual stimulus, the brain must maintain this information and avoid confounding it with other incoming visual signals, such as information about the location of a distracting visual stimulus (Katsuki and Constantinidis, 2012; Suzuki and Gottlieb, 2013). The mechanism that grants working memory robustness to such disturbances is not well understood (Lorenc et al., 2021; Wang, 2021).

Delay period activity in the dorso-lateral prefrontal cortex (dlPFC) has been linked to working memory related computations (Fuster and Alexander, 1971; Goldman-Rakic, 1995; Fuster, 2015, 2022; Constantinidis et al., 2018). These signals may represent the content of working memory or ongoing attentional and executive control processes (Lebedev et al., 2004; Sreenivasan et al., 2014; Lara and Wallis, 2015), and are central to perform working memory tasks (Bauer and Fuster, 1976; Funahashi et al., 1993; Buckley et al., 2009). The rich and heterogeneous nature of delay period activity in dlPFC has motivated different models of working memory maintenance, including persistent activity (Wang, 2001; Constantinidis et al., 2018), dynamic representations (Lundqvist et al., 2018; Miller et al., 2018) and activity silent mechanisms (Mongillo et al., 2008; Stokes, 2015).

To guide behavior, working memory information must be maintained in the presence of perturbations such as distractors. Some theories propose that PFC can robustly and stably maintain working memory information in the presence of other evolving signals by dissociating dynamic and persistent working memory representations (Druckmann and Chklovskii, 2012; Murray et al., 2017). This dissociation might protect working memory information from interference from other incoming sensory inputs which might act as distractors (Parthasarathy et al., 2019). Many studies have shown that working memory behavior is robust to perturbations from sensory distractors (Katsuki and Constantinidis, 2012; Suzuki and Gottlieb, 2013; Parthasarathy et al., 2017; Cavanagh et al., 2018). Delay period activity in PFC is affected by distractors, but working memory related signals are found to be largely preserved in the neural population activity, which aligns with behavior (Parthasarathy et al., 2017, 2019; Cavanagh et al., 2018).

Previous studies have sought to directly perturb delay period activity in dlPFC to influence memory-guided behavior. Long-lasting activity manipulations, e.g. by microstimulating dlPFC throughout the entire delay period (Opris et al., 2005a) or pharmacologically inactivating dlPFC (Sawaguchi and Iba, 2001), affect memory-guided behavior. However, transient optogenetic inactivation of PFC minimally impacts memory-guided behavior in monkeys (Mendoza-Halliday et al., 2023). Relatedly, transient optogenetic inactivation of premotor cortex in mice has no impact on prepared movements (Li et al., 2016; Inagaki et al., 2019). These studies also found that task-relevant signals in the delay period activity were not affected or quickly recovered from such transient manipulations.

Here, we tested whether dlPFC can robustly maintain memory-guided behavior under a diverse set of electrical microstimulation perturbations. We implanted monkeys with multi-electrode arrays in the dlPFC, which allowed us to transiently stimulate the area with a variety of microstimulation spatial patterns while they performed a memory-guided saccade task. We then simultaneously recorded the effect of microstimulation on dozens of neurons in the dlPFC. We found that patterned microstimulation broadly affected dlPFC neural population activity and caused strong changes in working memory-related signals. However, the monkeys’ behavior was minimally impacted. These observations could be reconciled when characterizing the effect of microstimulation at the population level. Microstimulation impacted activity in dimensions that reflected the natural time course of working memory representations. However, microstimulation minimally impacted dimensions that contained stable working memory signals, and activity in these dimensions quickly recovered from the perturbation. Our findings indicate that working memory signals in dlPFC are robust to a wide range of microstimulation perturbations, making memory-guided behavior robust to direct perturbations of brain activity.

## Materials and Methods

### Subjects and surgical procedures

We implanted a 96-electrode “Utah” Array (Blackrock Microsystems, Salt Lake City, UT) in the dlPFC of two adult, male rhesus macaques (*Macaca mulatta*) using sterile surgical techniques under isoflurane anesthesia. We implanted one array in the left 8Ar for Monkey W, and dual arrays in the left and right 8Ar for Monkey S (on the prearcuate gyrus, immediately anterior to the arcuate sulcus). The head was immobilized for recordings with a titanium headpost attached to the skull with titanium screws, implanted in a separate procedure prior to the array implants. Experimental procedures were approved by the Institutional Animal Care and Use Committee (IACUC) of Carnegie Mellon University and complied with guidelines set forth in the National Institute of Health’s Guide for the Care and Use of Laboratory Animals.

### Behavioral task

In each experimental session, the monkeys performed a memory-guided saccade task. On each trial, the monkeys first fixated on a dot at the center of the screen. After establishing fixation (for 100ms, monkey W; for 200ms, monkey S), one of four peripheral targets (45°, 135°, 225°, 315°) appeared on the screen for a brief period of time (100 ms, monkey W; 200 ms, monkey S). This was followed by a delay period, after which the center dot turned off (go cue) and the monkeys performed a saccade to the remembered target location to receive a liquid reward. In monkey W, the delay period was either 1.25 or 1.55 seconds in duration (with probabilities 0.8 and 0.2). In monkey S, it was 1.5 or 2 seconds in duration (with probabilities 0.5 and 0.5). The delay had different lengths so that the monkeys could not anticipate the go cue timing with certainty. Upon initiation of the saccade, the monkey’s eye position had to reach the peripheral target location within 200 ms and maintain gaze within 2.1° (monkey W) or 2.4° (monkey S) of the target center for 150 ms to receive a liquid reward. Stimuli were displayed on a 21” cathode ray tube monitor with a resolution of 1024×768 pixels and a refresh rate of 100 Hz at a viewing distance of 59 cm. Stimuli were generated using custom software written in MATLAB (MathWorks, Natick, MA) with the Psychophysics Toolbox extensions (Brainard 1997; Kleiner et al. 2007; Pelli 1997). Eye position was tracked monocularly using an infrared system at 1,000-Hz resolution (EyeLink 1000, SR Research, Mississauga, ON, Canada).

### Microstimulation experiments

We electrically microstimulated dlPFC on a subset of the trials in each session while the monkeys performed the memory-guided saccade task. On the rest of the trials, the monkey performed the task without microstimulation, which served as a control. On each trial, we stimulated on different electrodes of the array using single electrodes in monkey W and pairs of electrodes simultaneously in monkey S. By choosing different electrodes, we changed the spatial location of the stimulation on each trial, so we refer to each stimulation condition as a different “microstimulation pattern”. In each session, we stimulated with 3 (monkey W) or 4 (monkey S) different microstimulation patterns. We chose each pattern at random, but in some sessions we repeated some of the patterns used in previous days. Across sessions, we applied a total of 21 unique microstimulation patterns in monkey W and 4 unique microstimulation patterns in monkey S. Each session was organized in blocks. In each block, the experimental system performed for each trial a pseudorandomized selection of the different microstimulation conditions (3 or 4 microstimulation patterns, and no microstimulation) and the target angle conditions (4 target angles). We ran several blocks per session to ensure sufficient amounts of trials were collected under each target angle and microstimulation condition (about 20 trials per condition in monkey W and 30 trials per condition in monkey S). We refer to the set of trials collected under each microstimulation pattern as a “microstimulation experiment”. This includes trials across all target angle conditions. We performed 30 microstimulation experiments over the course of 10 sessions in monkey W (3 microstimulation patterns per session, 4 target angle angle conditions) and 4 microstimulation experiments over the course of 1 session in monkey S (4 microstimulation patterns, 4 target angle angle conditions).

### Microstimulation parameters

We used a Grapevine system and xippmex software to control microstimulation delivered using nano2+stim headstages (Ripple, Salt Lake City, UT). The microstimulation consisted of a 150 ms pulse train of 0.25 ms biphasic square pulses, with a frequency of 350 Hz and current of 50 or 125 μA per stimulated electrode. We kept these parameter values fixed for all sessions. The stimulation parameters were chosen based on previous microstimulation studies in dlPFC (Wegener et al., 2008), and we set the current amplitude low enough not to induce any eye movements. In monkey W, we stimulated on single electrodes with lower currents (50μA) because larger currents led to a substantial increase in trials in which fixation was aborted (though not consistent saccades). In monkey S, we stimulated with higher currents, using pairs of electrodes each at 125 μA. The reason we simulated at higher currents in monkey S was to induce sufficient activity modulation on the contralateral array. In this monkey, we could evoke saccades with some electrodes at microstimulation currents >300 μA. In monkey W, we stimulated using the array implanted in its left dlPFC and measured the effect of microstimulation on the same array. In monkey S, we stimulated only using the right dlPFC array and measured the effect on both the right and the left dlPFC arrays. In monkey W, the microstimulation was applied during the delay period at 500 ms after peripheral target offset. Even though the monkey could predict the time of the microstimulation, we stimulated at a different location in the array (or did not stimulate) randomly on each trial. In monkey S, we randomized the stimulation onset time (500, 600, or 700 ms after peripheral target offset).

### Neural recordings

Electrophysiological recordings were performed with a Grapevine acquisition system (Ripple, Salt Lake City, UT). Extracellular activity was recorded from the array, band-pass filtered (0.3–7,500 Hz), digitized at 30 kHz, and amplified by the system. Waveforms that exceeded a threshold were saved and stored for offline waveform classification. Thresholds were set by taking a multiple (4-5) of the root mean squared noise of the voltage measured on each channel. Waveforms were automatically classified as either noise or spikes using an artificial neural network (Issar et al., 2020). All spiking waveforms which survived this classification on a given channel were grouped together and treated as multi-unit activity.

### Data preprocessing

We excluded channels that in a given session had low spiking activity (firing rate < 1 sp/s) and high fano factors (> 8). To compute those criteria, we used activity during no-microstimulation trials. We also removed channels with a high percentage of coincident spikes, which is an indication that they could be electrically shorted (pairs with >20% coincident spiking within all 0.5 ms windows over the entire session; we removed one channel only from each electrically shorted pair). In monkey W, we included all channels from the left dlPFC array that passed the criteria (50-70 channels per session). In monkey S, we pooled the channels across the left and the right dlPFC arrays that passed the criteria (50 channels in the left array and 37 channels in the right array). We combined recordings across hemispheres because the recordings were from matched regions of cortex across the hemispheres and individual channels in each hemisphere showed microstimulation effects. Analysis of each hemisphere separately showed qualitatively similar results. Spike counts were binned in non-overlapping 50 ms windows. In our analysis of both microstimulation and no-microstimulation experiments, we focus on the period −300 ms before to 750 ms after microstimulation onset. To avoid microstimulation artifacts, we did not analyze activity from microstimulation onset to 50 ms after microstimulation offset. Given that the delay periods could have different lengths on each trial, the end of our analysis period (which we mark as t_end_ in our analyses) did not always correspond to the end of the trial (at go cue). However, since the delay times were randomized, the monkeys would not know whether the end of the trial would occur by the time the 750 ms mark was reached. In monkey S, given that we introduced variability in the microstimulation onset times, the period from microstimulation offset to go cue was shorter than 750 ms in some trials. This resulted in less trials contributing to some analyses for certain time periods after microstimulation, but we had a minimum of 16 trials per time step.

### Multi-unit analysis

We computed a measure of target angle selectivity to estimate the tuning strength of multi-unit activity, measured from each individual channel, at various times during the delay period (Fig. 1C,D and Fig. 2A,C). It was computed as the difference between the maximum and minimum firing rate across the 4 target angles at a given point in time (e.g. at t_post_). This measure can account for relative changes in tuning strength between target angle conditions, but it is not sensitive to changes in direction preference. We chose this measure given that the most prominent effect of microstimulation involved strong changes in modulation strength. We did not observe systematic changes in direction preference with microstimulation across units, although we did observe changes in direction preference in some units.

**Figure 1.**
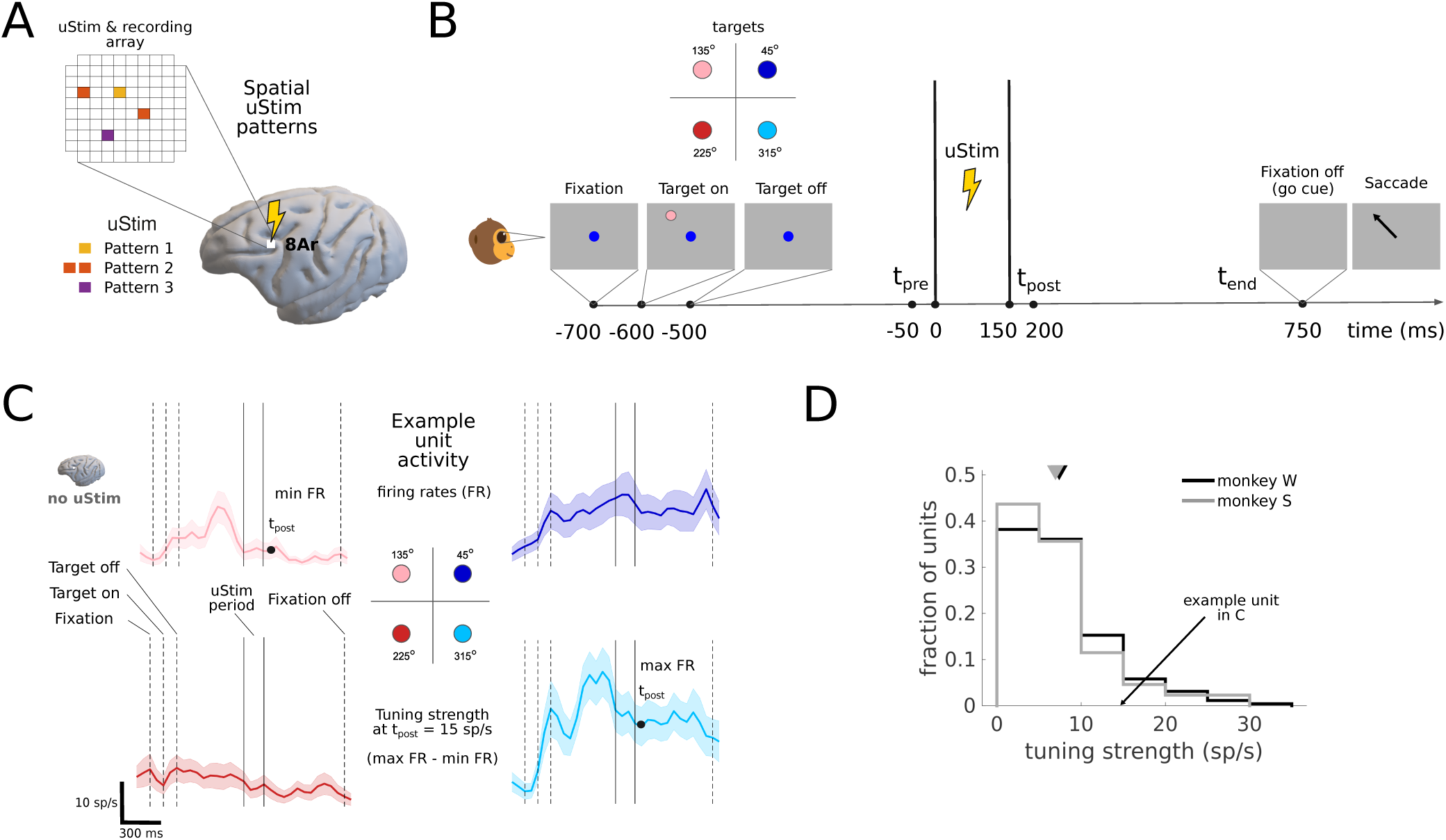
Multi-electrode electrical microstimulation to probe dlPFC delay period activity during working memory. **A**, Electrical microstimulation (uStim) protocol. An Utah array was implanted in the dlPFC, area 8Ar. Subthreshold stimulation was applied to either individual electrodes or pairs of electrodes in the array, creating a variety of spatial microstimulation patterns (see Methods). Neural population activity was simultaneously recorded from all channels in the array after microstimulation. **B**, Memory-guided saccade task. After the monkey acquired fixation, a target was briefly presented (for 100ms) at one out of four possible spatial locations. This was followed by a variable delay period (1.25-2s, see Methods), after which the goe cue signaled the monkeys to saccade to the remembered target location. Microstimulation was applied during the delay period, using 3-4 different spatial patterns on each session (see Methods). t_pre_: 50 ms before uStim onset; t_post_: 50 ms after uStim ends; t_end_: time of go cue. **C**, Neural activity of one representative unit for each of the four target angle conditions. Shown are firing rates (FR) computed from no-microstimulation trials (n=22) in an example session. This unit is spatially tuned to the different target locations during the delay period (activity is stronger for the rightward targets, and in particular for the 315° target, than for the other targets). Tuning strength at t_post_ = 15 spikes per second (sp/s), computed as max FR (target 315°) - min FR (target 225°). In no-microstimulation trials, t_post_ marks the same time as in microstimulation trials, but no stimulation was delivered in this case. Responses have been smoothed for visualization with a Gaussian filter with a standard deviation of 40 ms. Monkey W, session 20220810, unit 30. **D**, Tuning strengths across all units at t_post_ (monkey W, n=10 sessions; monkey S, n=1 session). Triangles indicate mean tuning strength across all units and sessions (7±6 sp/s, mean±std, monkey W; 7±6 sp/s, monkey S).

**Figure 2.**
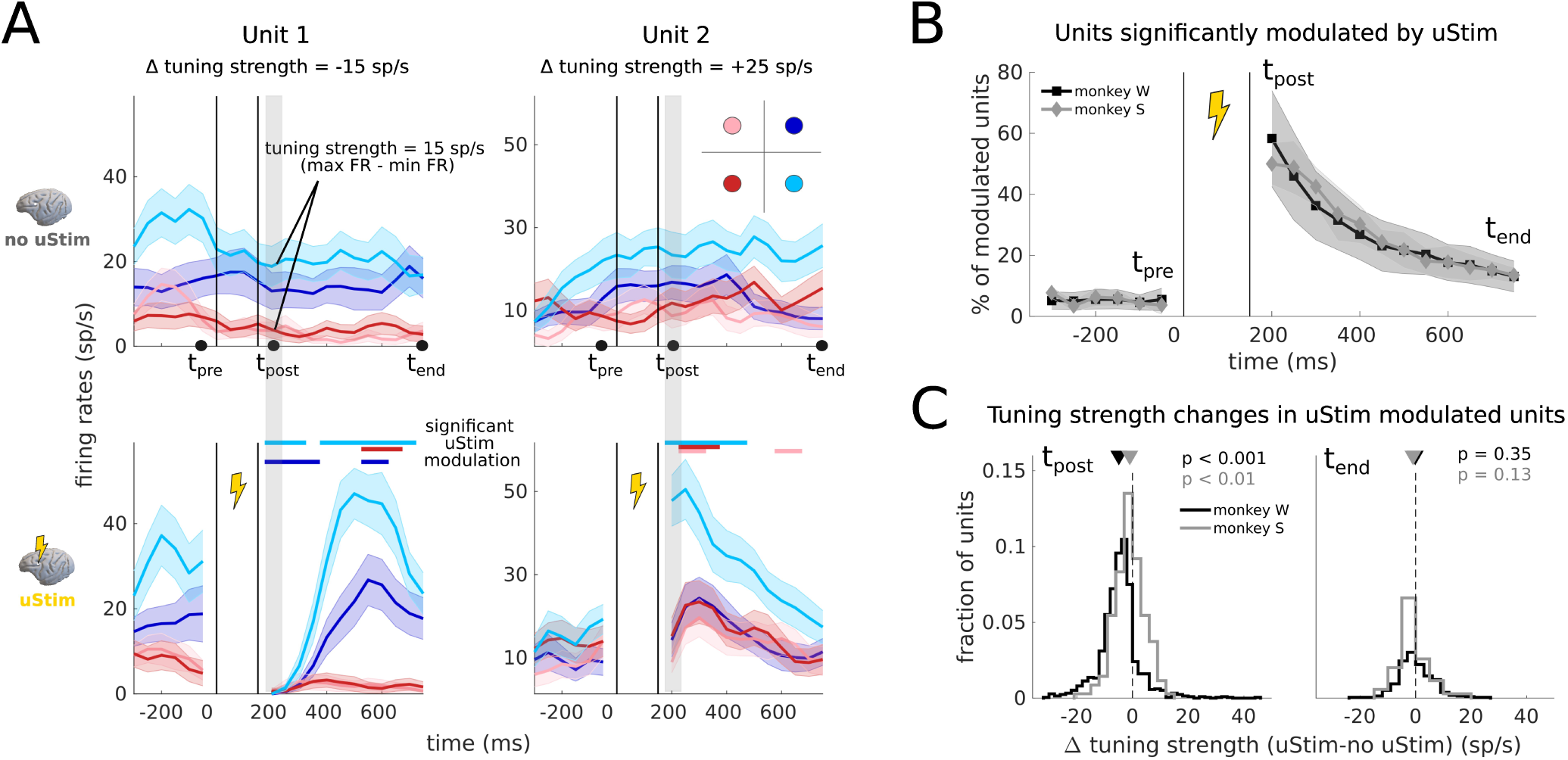
Microstimulation impacts working memory related signals in dlPFC. **A**, Effect of microstimulation on neural activity and target angle tuning for two example units (left panel, same unit as in Fig. 1C). Top, PSTHs for trials without microstimulation (no-uStim). Bottom, PSTHs for trials with microstimulation (uStim) Microstimulation can induce both decreases (left panels) or increases (right panels) in activity. Furthermore, the tuning strength can decrease (−15 sp/s at t_post_, left panels) or increase (+25 sp/s at t_post_, right panels). Colored bars above the bottom panels indicate significant microstimulation-induced activity modulation for each target angle compared to no-microstimulation activity for the same target angle (p<0.05, Wilcoxon rank sum test, n=22 trials per condition). Responses have been smoothed for visualization with a Gaussian filter with a standard deviation of 40 ms. Monkey W, session 20220810, units 30 and 56. **B**, Percentage of units that are significantly modulated by microstimulation at different times during the delay period (mean±std across all microstimulation experiments; monkey W, n=30; monkey S, n=4; p<0.05, Wilcoxon rank sum test). **C**, Tuning strength changes induced by microstimulation at t_post_ and t_end_ in units significantly modulated by microstimulation. Tuning strength was predominantly reduced by microstimulation at t_post_ (p<0.001, monkey W; p<0.01, monkey S; one-tailed t-test; all units across all microstimulation experiments). Triangles indicate mean tuning strength across all units and sessions for each monkey. By t_end_, there was no longer a significant decrease in tuning strength on average across units (p=0.35, monkey W; p=0.13, monkey S; one-tailed t-test), although some units remained affected.

### Behavioral analysis

Saccade precision was estimated as the distance between the saccade endpoint to the target (in degrees of visual angle) (Fig. 3B). Reaction times were computed as the time between go cue (i.e. fixation dot offset) and the time the monkey’s eye crossed the boundary of the virtual fixation window (Monkey W, 1.4° in diameter around the fixation point; Monkey S, 2.7°) (Fig. 3C). To compute the fraction of target misses (target misses/(target misses + correct trials), Fig. 3D), we considered all trials in which the monkey maintained fixation throughout the delay until the go cue. We excluded fixation breaks where the monkey left the fixation window before the go cue. In correct trials, the monkey reached the target window and maintained their eyes within that window for 150 ms. All other trials constituted “target misses”, which included cases where monkeys initiated a saccade toward the target location but failed to reach the target window (which was the vast majority of the errors), as well as a very small number of cases where the monkey failed to initiate a saccade or performed a saccade toward an incorrect location. The monkeys were able to correctly report the memorized target location in this task equally well with and without microstimulation. Because of this, we focused primarily on analyzing neuronal activity from correct trials.

**Figure 3.**
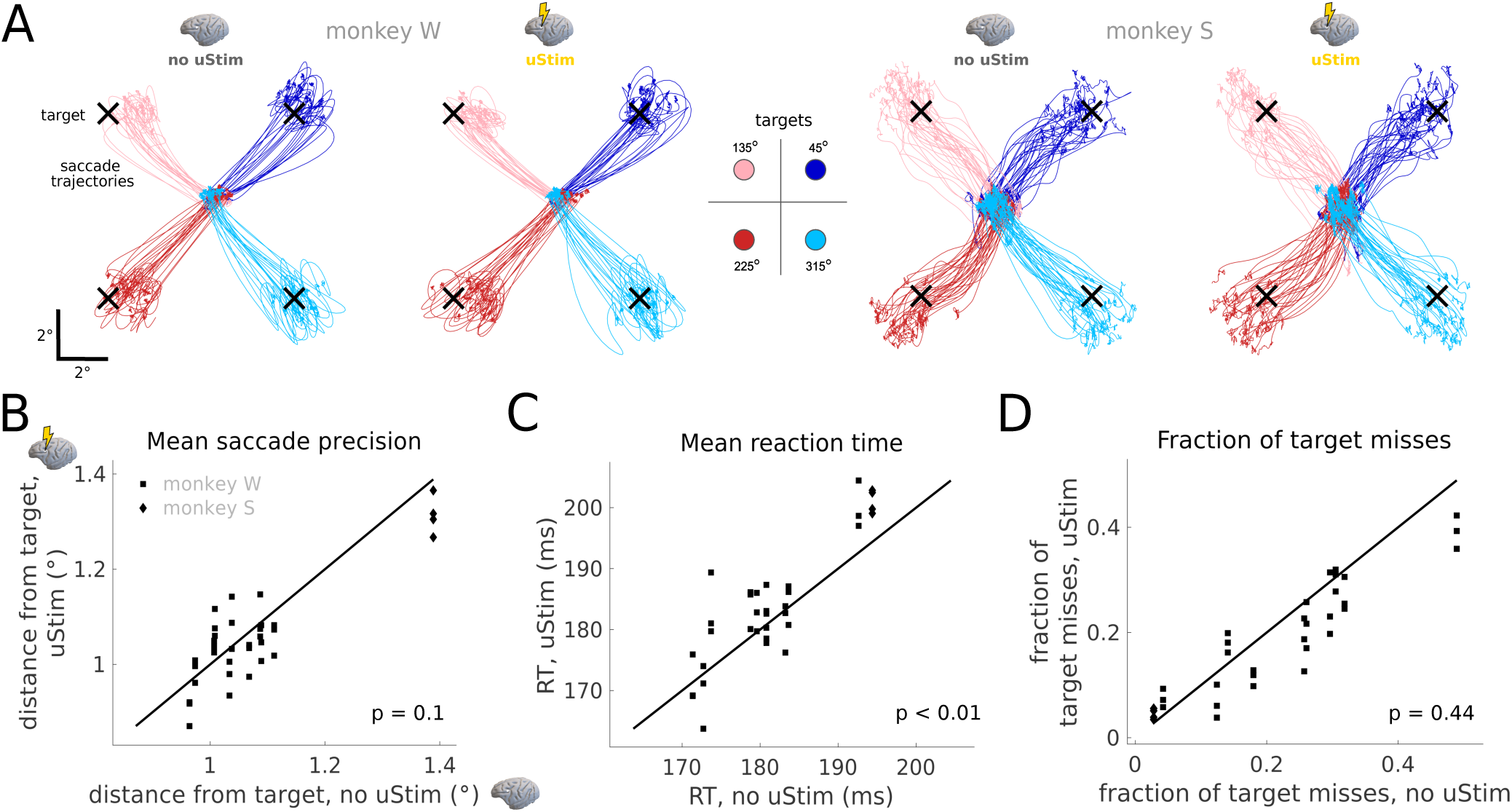
Microstimulation in dlPFC minimally impacts memory-guided behavior. **A**, Saccade trajectories in microstimulation (uStim) and no-microstimulation (no-uStim) trials for an example session of each monkey. Monkey W, session 20220801; monkey S, session 20210311. Trajectories shown from go cue to target acquisition, for all target angle conditions (trials per condition, n=22, monkey W; n=33, monkey S). Trajectories have been smoothed for visualization with a Gaussian filter with a standard deviation of 2 ms. **B**, Mean saccade precision in microstimulation versus no-microstimulation trials, measured as the trial-averaged distance from the saccade endpoint to the target. Distances on microstimulation trials are not significantly different from distances on no-microstimulation trials (p=0.1, two-tailed t-test; all microstimulation experiments across both monkeys, n=34). **C**, Mean reaction time (RT) on microstimulation versus no-microstimulation trials. RTs on microstimulation trials are significantly higher than on no-microstimulation trials (p<0.01, two-tailed t-test; n=34), but only by 3 ms on average. **D**, Fraction of target misses (see Methods) on microstimulation versus no-microstimulation trials. The fraction of target misses on microstimulation trials is not significantly different than on no-microstimulation trials (p=0.44, two-tailed Wilcoxon rank sum test; n=34). Each individual square/diamond in panels B-D indicates a different microstimulation experiment (which includes all trials for a given microstimulation pattern within a session, considering all target angle conditions). The squares/diamonds appear organized in columns along the horizontal axis because each microstimulation experiment within a session was compared against the same no-microstimulation experiment (one per session).

### Classification analysis

We trained a Poisson Naive Bayes classifier to predict target angle identity on single trials at different time points during the delay period (Fig. 4). We trained the classifier separately for microstimulation and no-microstimulation experiments. A different classifier was trained for each microstimulation experiment. We used leave-one-trial-out cross-validation to train and test the classifier, and used activity from all units. A different classifier was trained independently at each time point during the delay period.

**Figure 4.**
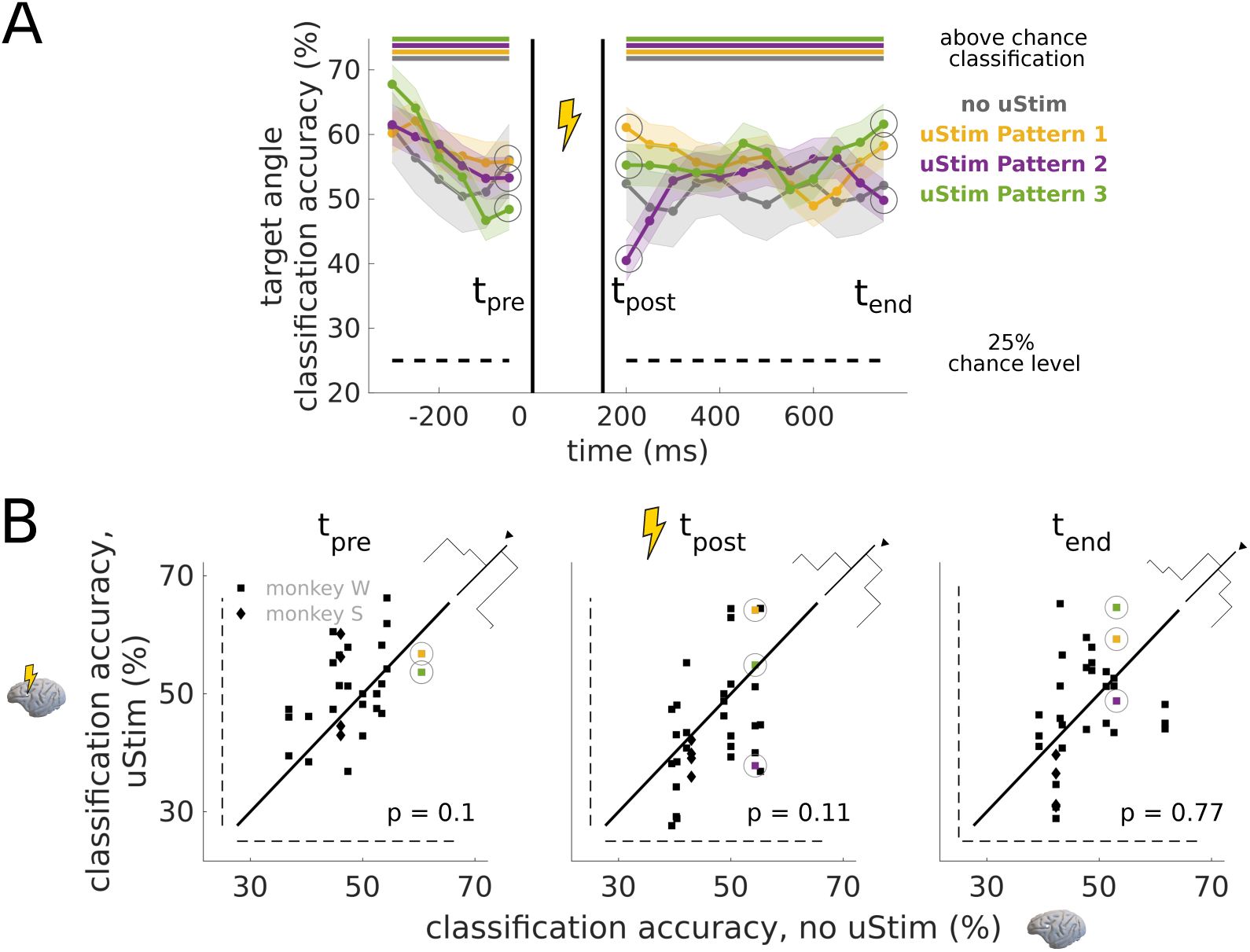
Working memory-related information in dlPFC is preserved after microstimulation. **A**, Cross-validated target angle classification accuracy for microstimulation (uStim) and no-microstimulation (no-uStim) experiments. Example session with three microstimulation experiments (colored lines) and one no-microstimulation experiment (gray line). A separate classifier was trained on each time step for each experiment. Classification was well above chance (dashed line, chance probability: 0.25) throughout the delay period, even at t_post_ (colored bars; p<0.05, binomial test). Classification accuracies were similar in microstimulation and no-microstimulation experiments. Lines have been smoothed for visualization with a Gaussian filter with a standard deviation of 40 ms. Monkey W, session 20220802. **B**, Classification accuracies for all microstimulation experiments at different times during the delay period. Classification accuracy was not significantly different in microstimulation versus no-microstimulation experiments (left to right panels for t_pre_, t_post_ and t_end_; p=0.1, p=0.11 and p=0.77, two-tailed Wilcoxon rank sum test; n=34). In all cases, classification was above chance (dashed lines), even at t_post_ (middle panel). Gray circles and colored squares indicate the example experiments shown in panel A, but here with unsmoothed classification accuracies. The purple square in the left panel overlaps with the green square and is therefore not visible.

To predict target angle identity based on activity within the memory subspace alone we used a different classification procedure. First, we projected the activity of no-microstimulation trials into the memory subspace and computed class means across trials and time for each target angle condition. Second, we projected the activity of microstimulation trials onto the memory subspace and classified each trial based on the euclidean distance to the different class means. Even though class means were computed from no-microstimulation trials, and that the means were based on averages across time, we could accurately and stably classify microstimulation trials at various times during the delay period (see last section of Results).

### Population analysis

We used two different dimensionality reduction methods, Factor analysis (FA) and demixed PCA (dPCA) to estimate different subspaces of the neural population activity. FA is a statistical method whose objective is to find the “dominant” dimensions that capture the greatest shared variance among the neural units. We sought to estimate the dominant dimensions of the neural activity under natural task conditions, so we fit FA to no-microstimulation trials exclusively. In each session, we fit FA to the binned spike counts (50 ms bins) from each unit during the delay period in all no-stimulation trials (including all target angle conditions). The dimensionality of the “dominant subspace” (4D) was estimated by computing the optimal dimensionality separately for each session based on cross-validated data likelihoods (Santhanam et al., 2009), averaging the estimates across all sessions, and taking the rounded average. Low-dimensional trajectories were inferred by the model based on posterior mean estimates.

To find the “memory subspace”, we used demixed dPCA (Kobak et al., 2016). dPCA’s objective is to find dimensions of the neural activity that are predominantly related to specific variables. We used dPCA to find memory dimensions with target angle variance that were immune to the effects of the microstimulation, and that contained target angle signals that were stable over time. To do this, the dPCA model was fit to both microstimulation and no-microstimulation trials, and different subspaces were found that “demixed” target angle, microstimulation and time related signals in the population. We refer to the target angle dimensions as the “memory subspace”. We fitted dPCA to the firing rates from each unit estimated in 50 ms bins during the delay period, averaged across trials of a given target angle and microstimulation condition. We set the dimensionality of the memory subspace to be 4D, to match the dimensionality of the dominant subspace. This was necessary to be able to fairly compare activity across the two subspaces. We confirmed that the four “memory” dimensions explained on average 75% of the target angle variance in the averaged neural activity across sessions. This subspace contained only a small fraction of time-related and stimulation-related variance, and thus, target angle signals in this subspace were largely stable over time and were minimally influenced by microstimulation.

The dPCA objective explicitly decorrelated (or “demixed”) dimensions with target angle variance from dimensions with microstimulation variance, as well as from dimensions with time related variance. The FA model, on the contrary, did not include any objective of this type. FA’s goal was to capture the dominant modes of co-fluctuation of the population activity, and not necessarily variance due to specific task variables. However, the neural population during natural task conditions was driven by co-variations of activity between units across target angle conditions, as well as time. This is why we observed such signals in this subspace (Fig. 5A). In contrast, it was not necessarily a given that these natural modes of the activity would be modulated by the microstimulation, as we indeed observed (Fig. 5A).

**Figure 5.**
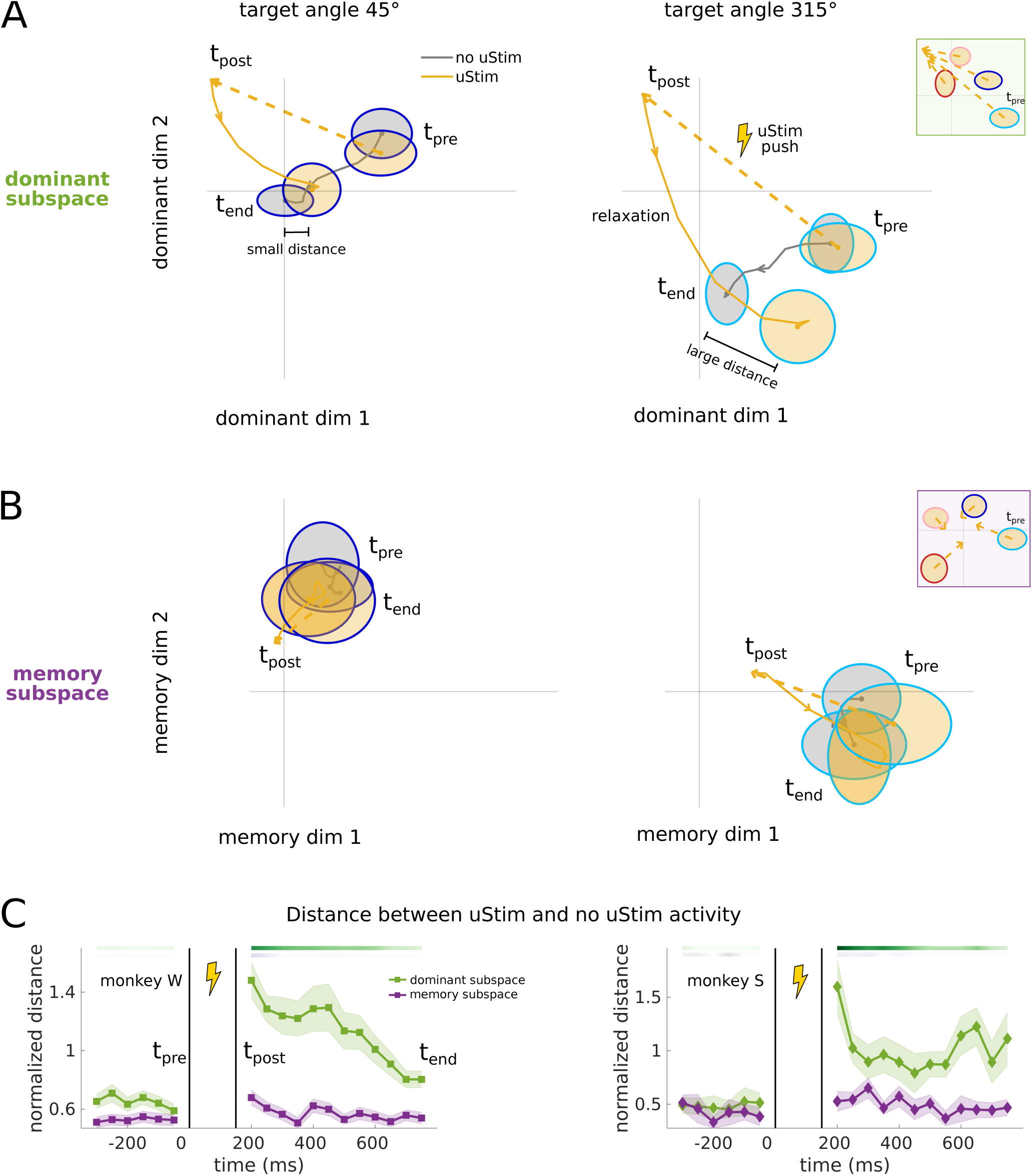
Microstimulation minimally impacts activity in a memory subspace of dlPFC neural population activity. **A**, Averaged neural trajectories in the dominant subspace during example microstimulation (uStim, yellow lines) and no-microstimulation (no-uStim, gray lines) trials, for two example target angle conditions (45° and 315°, left and right panels). Trajectories are shown from t_pre_ to t_end_ (except for the period t_pre_ to t_post_ during microstimulation, where data were excluded because of microstimulation artifacts; yellow dashed arrows indicate this period, illustrating the change in activity induced by microstimulation). Inset at the top right corner shows activity at t_pre_ and t_post_ during microstimulation trials for all target angle conditions. 95% confidence intervals (CI) across all trials (n=22) are shown at t_pre_ and t_end_. Microstimulation strongly modulates activity in the dominant subspace (yellow dashed arrows are long), and the activity does not always recover to its natural state by t_end_ (distance between microstimulation and no-microstimulation activity distributions at t_end_). Monkey W, session 20220810. **B**, Averaged neural trajectories in the memory subspace. Same conventions as in panel A. Microstimulation modulation is weak in this subspace (yellow dashed arrows are short), and the activity remains in the same location throughout the delay period. Trajectories in panels A and B have been smoothed with a Gaussian filter with a standard deviation of 60 ms. Monkey W, session 20220810. **C**, Mean distance between microstimulation and no-microstimulation activity throughout the delay period in the dominant and memory subspaces (green and purple lines; mean±CI across all microstimulation conditions; distances are normalized to the standard deviation of the activity in the no-microstimulation condition of each session). Color bars indicate percent of conditions with significant microstimulation-induced activity modulation in the dominant (top green bar) and memory (bottom purple bar) subspaces (p<0.05, two-tailed multivariate test). Here a “condition” refers to a target angle condition of a microstimulation experiment (monkey W, n=120; monkey S, n=16). Color scale goes from 80% (darkest) to 0% (brightest) in monkey W. In monkey S, the color scale goes from 100% to 0%. In panels A and B, we show population activity in the two leading dimensions of the dominant and the memory subspaces. The quantification in panel C was computed using all four dimensions of each subspace.

The dominant and memory subspaces overlapped along some dimensions (minimum subspace angle = 55°), and both contained signals related to working memory, meaning that target angle classification could also be performed in the dominant subspace. However, we found that classification accuracies were lower in the dominant subspace than in the memory subspace.

The distance metric in Fig. 5C was computed as follows: First, we calculated the difference between the mean activity in microstimulation and no-microstimulation conditions along each dimension in the subspace (4 dimensions). Second, we standardized the differences by the standard deviation of the activity in no-microstimulation conditions along each dimension. Third, we computed the vector norm of the differences across all dimensions. In this way, we could compare distances across sessions and monkeys.

### Statistical tests

To test whether microstimulation and no-microstimulation activity samples were statistically different, we used a two-tailed two-sample Wilcoxon rank sum test at 5% significance level (Fig. 2A). In Fig. 2B, we applied this test considering activity across all four target angle conditions. To test for significance of microstimulation-induced decreases in tuning strength (Fig. 2C), we used a one-tailed two-sample t-test. To test for significance of microstimulation-induced changes in accuracy (Fig. 3B) and reaction times (Fig. 3C), we used a two-tailed two-sample t-test. To test for microstimulation-induced changes in target misses (Fig. 3D) and classification accuracies (Fig. 4B), we used a two-tailed two-sample Wilcoxon rank sum test. To assess whether we could classify above chance, we used a binomial test at 5% significance level to compare the obtained classification probabilities against the 0.25 chance level probability (given four possible classes, the four target angle conditions). To compare microstimulation and no-microstimulation activity within the dominant and memory subspaces (Fig. 5C), we used a multivariate two-sample test at 5% significance level based on the statistical energy of the samples (Aslan and Zech, 2005), which compares the two sample distributions in high-d.

## Results

We designed an experimental protocol using electrical microstimulation to probe delay period activity in dlPFC during working memory. We implanted two monkeys with multi-electrode arrays in dlPFC, area 8Ar (Fig. 1A), and applied different microstimulation (uStim) patterns while they performed a memory-guided saccade task (Fig. 1B). We used microstimulation patterns below the threshold that evoked saccades, consisting of either single electrodes or pairs of electrodes at different spatial locations in the array (Fig. 1A, Methods). In area 8Ar, the current thresholds required to evoke saccades tend to be higher than the nearby frontal eye fields (Bruce et al., 1985), though subthreshold stimulation can still impact behavior (Opris et al., 2005a). We focused on area 8Ar in the dlPFC since it is implicated in spatial working memory and contains rich and heterogenous signals related to the transformation of sensory inputs to oculomotor actions in memory-guided saccade tasks (Funahashi et al., 1989; Opris et al., 2005a; Leavitt et al., 2018; Khanna et al., 2020).

At the beginning of the task, a target was briefly shown at one out of four possible spatial locations (Fig. 1B, Methods). This was followed by a variable delay period (1.5-2s; from “target off” to “fixation off” in Fig. 1B), after which the animal reported the location of the target with a saccadic eye movement to the remembered location. We stimulated during the delay period of the task when only a fixation spot was present on the screen and the animal had to hold in memory the location of the presented target. On each trial, we stimulated with a different microstimulation pattern (Fig. 1A), with 3-4 patterns selected for each experimental session (Methods). Interleaved with the microstimulation trials, we also ran trials where no microstimulation was applied. We consider a “microstimulation experiment” as the collection of trials collected under a given microstimulation pattern within a session, which includes trials for all target angle conditions (Methods). Similarly, we consider a “no-microstimulation experiment” as the collection of trials collected with no-microstimulation in a given session, for all target angle conditions.

Recordings from different channels in the array revealed delay period activity that was tuned to the different spatial locations (Fig. 1C, example unit), as previously reported in this area (Funahashi et al., 1989; Leavitt et al., 2018; Khanna et al., 2020). Some of these signals were persistent throughout the delay (e.g. target 45 activity, Fig. 1C), a canonical feature of delay period activity in the prefrontal cortex (Fuster and Alexander, 1971; Constantinidis et al., 2018). Other neuronal responses were dynamic (e.g. target 135 activity, Fig. 1C), consistent with more recent descriptions of working memory signals in PFC (Murray et al., 2017; Lundqvist et al., 2018; Khanna et al., 2020; Wang, 2021). Across units and sessions, tuning strength (a measure of selectivity to different targets, Methods) ranged from a few spikes per second (sp/s) to over 30 sp/s (Fig. 1D). Thus, the dlPFC neural population contained working memory related signals with target angle location information of the type needed to solve the task.

### Microstimulation broadly affected dlPFC delay period activity during working memory

Having identified working memory related signals in the dlPFC population during the delay period, we sought to understand how different microstimulation perturbations impacted these signals in individual neurons. For this, we measured the effect of microstimulation on the activity across all recorded units in the array and quantified the impact on tuning strength.

We found that microstimulation disrupted working memory related signals in dlPFC. Neural activity was strongly modulated by microstimulation, and this modulation induced strong changes in tuning strength, even in correct trials (Fig. 2; example units). We observed that microstimulation often strongly suppressed activity (Fig. 2A, top panel, no microstimulation; bottom panel, microstimulation). This suppression happened in all target angle conditions for many units, resulting in strong reductions of tuning strength (e.g. from 15 sp/s to 0 sp/s at microstimulation offset, t_post_; Fig. 2A, top vs. bottom left panels). In some cases, microstimulation also caused increases in activity for some conditions (Fig. 2A, top vs. bottom right panels) and concomitant increases of tuning strength (e.g. from 15 sp/s to 40 sp/s at t_post_; Fig. 2A, top vs. bottom right panels).

The microstimulation effect on neural activity was widespread across the dlPFC population, with over 50% of the recorded units significantly modulated during one or more of the four target angle conditions immediately after microstimulation offset (at t_post_, Fig. 2B; monkey W: 58±16, mean±std, n=30 microstimulation experiments; monkey S: 50±7, mean±std, n=4 microstimulation experiments). Some units remained modulated by the end of the trial (at t_end_; Fig. 2B; monkey W: 13±5, mean±std, monkey S: 14±3, mean±std). This indicates that activity did not always recover from the microstimulation perturbation by the time the monkeys were instructed to perform the saccade.

Next, we sought to quantify the impact of microstimulation on working memory related signals in units that were significantly modulated by microstimulation. In many units, the microstimulation perturbation caused strong changes in tuning strength (Fig. 2C). The modulated units experienced both decreases and increases in tuning strength, but overall there was a significant reduction in tuning strength right after microstimulation offset (Fig. 2C, left panel). The changes in tuning strength were often substantial (>10 spikes per second). By the end of the trial, there was no longer a significant decrease in tuning strength on average across experiments (Fig. 2C, right panel), though some units remained affected. Thus, we found that microstimulation broadly impacted activity across the dlPFC neural population and produced strong and long lasting changes in tuning strength, affecting working memory related signals that may be crucial for the task.

### Microstimulation minimally impacted memory-guided behavior

Having found that there was a pronounced effect of microstimulation on neural activity (Fig. 2), we next sought to determine whether microstimulation impacted working memory behavior. We asked whether microstimulation had an impact on the eye movements to remembered targets. We observed that saccade trajectories were qualitatively similar in microstimulation versus no-microstimulation experiments (Fig. 3A). However, since microstimulation can have subtle impacts on behavior (Opris et al., 2005a, 2005b; Murphey and Maunsell, 2008), we quantified the impact of microstimulation on several eye movement metrics.

First, the precision of the saccades (distance of the saccade endpoint to the presented target) was not impacted by microstimulation. Across all microstimulation experiments in both animals, we found no significant difference in saccade precision between microstimulation and no-microstimulation experiments (Fig. 3B).

Second, we asked whether microstimulation had an impact on the readiness of the animals to report the remembered target location by examining the saccadic reaction time (RT, Fig. 3C). The overall RT across experiments was about 3 ms faster in the no-microstimulation versus microstimulation experiments, which was small compared to the typical range of RTs within a given experiment (no-microstimulation, 181±7 ms; microstimulation, 184±10 ms; mean±std). Nonetheless, this difference was statistically significant. This small but significant difference could indicate that the microstimulation had a slight tendency to disrupt the memory signal or the saccadic preparation (Opris et al., 2005a; Churchland and Shenoy, 2007).

Third, microstimulation did not increase the rate of errors (Fig. 3D). We considered target misses, which applied to cases where monkeys failed to report the correct target location after go cue presentation (see Methods). This included cases where monkeys initiated a saccade toward the target location but failed to reach the target window (which was the vast majority of the errors), as well as a small number of cases where the monkey failed to initiate a saccade or performed a saccade toward an incorrect location. We found no statistically significant difference in the fraction of target misses in microstimulation versus no-microstimulation trials (Fig. 3D, individual squares). We can thus conclude that the observed microstimulation-induced neural activity changes have little or no consequence for the behavioral performance of the monkeys.

### dlPFC preserved working memory related information after microstimulation

How can we reconcile the pronounced microstimulation effects on neural activity (Fig. 2) with the little or no impact of microstimulation on behavior (Fig. 3)? One possible explanation is that the area we are perturbing, despite containing signals relevant to the task, is not causally implicated in the generation of the behavior. However, the large body of literature involving lesions, cooling and inactivation in the dlPFC points at a critical and necessary role of this region in this type of working memory behavior (Bauer and Fuster, 1976; Funahashi et al., 1993; Buckley et al., 2009). An alternative explanation is that, similar to what has been previously reported with sensory distractors (Parthasarathy et al., 2017, 2019; Cavanagh et al., 2018), the induced neural changes from microstimulation do not entirely disrupt working memory information.

To test this, we analyzed the dlPFC neural population activity to determine if working memory-related information was preserved in the population after microstimulation. To extract this information from the neural population, we trained target angle classifiers using delay period activity separately for microstimulation and no-microstimulation experiments. Furthermore, a separate classifier was trained at each time step. We found that we could classify well above chance in both cases and with similar accuracy in both microstimulation and no-microstimulation experiments (Fig. 4A, example session with three different microstimulation experiments). Importantly, we could classify similarly well throughout the entire delay period. Across all experiments we performed, classification accuracy was not significantly different in microstimulation versus no-microstimulation experiments (Fig. 4B, shown at t_pre_, t_post_ and t_end_). There was variability in classification accuracy across microstimulation experiments (individual squares and diamonds), but in all cases we could classify above chance (dashed lines), even immediately after microstimulation (at t_post_, middle panel). This analysis demonstrates that dlPFC robustly encodes working memory related information after microstimulation. The presence of working memory related signals in the neuronal population activity may explain why behavior is not disrupted by perturbations that have strong effects on the activity of individual units.

### dlPFC’s neural activity was minimally impacted by microstimulation in a memory subspace of the neural population activity

Having found that working memory information is preserved in the dlPFC population after microstimulation, we next sought to find which dimensions of the population activity were perturbed by microstimulation. We considered two subspaces. First, we applied Factor Analysis (FA) to extract the dimensions of the activity that captured the greatest shared variance among the neurons under natural task conditions (i.e. during no-microstimulation) (Fig. 5A, the “dominant subspace”, see Methods). Second, we used demixed PCA (dPCA) (Kobak et al., 2016) to specifically look for dimensions of the activity that were not affected by microstimulation, and which could contain stable working memory related signals during the delay (Fig. 5B, the “memory subspace”, see Methods). The existence of these dimensions might explain the robustness of behavior, since working memory related signals could be stably read out throughout the delay (Parthasarathy et al., 2019).

We found that microstimulation strongly impacted activity in the dominant subspace (Fig. 5A). The effect of microstimulation could be visualized as a push of the neural activity along specific directions (dashed yellow arrows) and away from its natural time course (gray trajectories, no microstimulation). At microstimulation offset (t_post_), the activity relaxed toward the end location of the trajectories under natural task conditions (yellow trajectories; t_end_ in no microstimulation). In this dominant subspace, the effect of microstimulation was easily evident (dashed arrows), and the activity did not always recover to its natural state by the end of the trial (left panel, recovery, vs. right panel, non-recovery; t_end_ locations, 95% CI, n=20 trials). These population-level effects match the observations made at the individual unit level, where we found strong microstimulation modulation at t_post_ and a persistent modulatory effect until the end of the trial for a subset of the units (Fig. 2). By contrast, we found that microstimulation weakly affected activity in the memory subspace (Fig. 5B). Modulation between t_pre_ and t_post_ was small (length of dashed arrows) and the activity recovered to its natural state by the end of the trial (t_end_, no-microstimulation).

Consistently across microstimulation experiments, population activity was strongly modulated by microstimulation in the dominant subspace, whereas it was minimally impacted by microstimulation in the memory subspace (Fig. 5C). To see this, we quantified the distance between microstimulation and no-microstimulation activity at various times during the delay period. Activity after microstimulation strongly deviated from no-microstimulation activity in the dominant subspace, particularly at t_post_ (green lines and shades, mean±CI across experiments). This indicated that microstimulation strongly pushed activity away from its natural state. After this push, the activity did not always recover by the end of the trial (green bar, color scale: % of experiments with significant activity modulation due to microstimulation in the dominant subspace; monkey W, at t_post_: 74%, at t_end_: 15%, n=120 microstimulation and target angle conditions; monkey S, at t_post_: 100%, at t_end_: 19%, n=16). By contrast, activity in the memory subspace remained minimally impacted by microstimulation consistently across microstimulation experiments (purple lines and shades, mean±CI), and the modulation was largely absent by the end of the trial (purple bar, color scale: % of experiments with significant activity modulation due to microstimulation in the memory subspace; monkey W, 25% at t_post_, 1% at t_end_, n=120; monkey S, 12% at t_post_, 0% at t_end_, n=16).

The dominant subspace captured two prominent features of the population activity: working memory tuning and the time evolution of responses (Fig. 5A). The activity occupied different locations in the space depending on target angle condition (left vs. right panels, 45° and 315°; inset, all angles), reflecting target tuning. Additionally, neural activity during the delay naturally evolved from one location (at t_pre_) to a different location (at t_end_) (no-microstimulation experiments). The dominant subspace captured about 50% of the target angle variance in the population activity (monkey W: 54±13%, mean±std, n=10 sessions; monkey S: 57%, n=1; Methods), and 30 to 50% of time-related variance (monkey W: 51±19 %, n=10; monkey S: 28%, n=1) in both no-microstimulation and microstimulation conditions. These findings indicate that the dominant subspace contained working memory related signals that evolved over time.

Contrary to what was found in the dominant subspace (Fig. 5A), the activity in the memory subspace remained in roughly the same location throughout the entire delay period, in both microstimulation and no-microstimulation experiments. This location depended on the target angle condition (left vs. right panels, 45° and 315°; inset, all angles), indicating target tuning. The memory subspace captured about 75% of the target angle variance in the population activity in both no-microstimulation and microstimulation conditions (monkey W: 75±8%, mean±std, n=10 sessions; monkey S: 75%, n=1; Methods), and only about 2% of time related variance (monkey W: 3±1 %, n=10; monkey S: 2%, n=1). The small amount of time-related variance indicates that the memory subspace contained stable working memory information throughout the delay. This might explain the robustness of behavior after microstimulation, since working memory related signals could be stably read out with the same decoder throughout the delay.

To test this possibility, we attempted to extract working memory related information from this memory subspace before and after microstimulation. Using a static classification method based on no-microstimulation activity from the memory subspace alone (see Methods), we could stably and accurately read out target angle information throughout the delay period (classification accuracy during microstimulation: monkey W, t_pre_: 57±2%, t_post_: 48±3%, t_end_: 54±3%, mean±CI, n=30 microstimulation experiments; monkey S, t_pre_: 57±6%, t_post_: 39±5%, t_end_: 39±13%, mean±CI, n=4; above chance classification). It was also possible to read out working memory related information in the dominant subspace, albeit with lower accuracy (see Methods). Importantly, classification accuracies obtained with this static classification method were similar to those obtained using classifiers with time-varying parameters, and which relied on all dimensions of the data (Fig. 4). This meant that working memory related signals could be stably read out with the same decoder throughout the delay, making this information readily available to downstream areas (Parthasarathy et al., 2019).

In summary, we found that dynamic and stable working memory representations coexist in dlPFC, and that microstimulation differentially impacts them. In dimensions where working memory signals naturally evolved over time, microstimulation strongly modulated neural activity. In contrast, in dimensions containing working memory signals that were stable over time, microstimulation minimally impacted neural activity, and working memory information could be stably read out throughout the delay period.

## Discussion

Here, we studied working memory computations by directly perturbing delay period activity in the dlPFC using electrical microstimulation. We implanted monkeys with multi-electrode arrays in dlPFC, which allowed us to stimulate with a variety of spatial microstimulation patterns while monitoring the effect on the recorded neural population. We found that microstimulation broadly affected the activity of individual neurons in dlPFC, including changes to the tuning strength of individual neurons that displayed working memory-related activity. However, we found minimal impact on the ability of the animals to correctly perform the task. This apparent contradiction was reconciled at the population level, where we found that working memory information was preserved in dlPFC after microstimulation, and that the information could be stably read out from a specific “memory subspace” of the neural population activity. Our findings indicate that working memory exhibits robustness to direct microstimulation perturbations in the dlPFC.

PFC might be endowed with circuit properties that grant working memory signals stability to diverse activity perturbations. For example, a previous study also found a stable memory subspace that was not affected by perturbations, but in this case these were induced by sensory distractors (Parthasarathy et al., 2019). Cognitive-related signals that are not disrupted by external stimuli or contextual events have also been found in PFC during executive control functions other than working memory, such as categorical reasoning (Freedman et al., 2003; Cromer et al., 2011), rule-based decision-making (Mante et al., 2013; Siegel et al., 2015) and selective attention (Snyder et al., 2021). This feature might be an intrinsic and general property of high order areas such as PFC, which is required to form and maintain stable cognitive states to guide behavior (Snyder et al., 2021).

PFC might robustly and stably maintain working memory information in the presence of other evolving signals by representing stable and dynamic information in different subspaces of the neural population activity (Druckmann and Chklovskii, 2012; Murray et al., 2017). We found such subspace dissociation of stable and dynamic working memory signals in the dlPFC population (Fig. 6A, signals in “memory” versus “dominant” subspaces). Importantly, while the “stable” memory subspace was minimally affected by microstimulation, the “dynamic” dominant subspace was strongly modulated by microstimulation (Fig. 6B). The dissociation of stable and dynamic variables in different subspaces of dlPFC’s neural activity might underlie working memory robustness to sensory perturbations (Parthasarathy et al., 2019), as well as to microstimulation perturbations. A similar subspace dissociation has also been proposed in the motor cortex as a mechanism to separate preparatory activity from movement activity so that movements are not prematurely generated during motor planning (Kaufman et al., 2014; Elsayed et al., 2016). Related mechanisms might be in place to prevent cognitive and arousal-related signals from influencing motor responses (Johnston and Smith, 2024), or value-related signals from prematurely driving choice (Yoo and Hayden, 2020).

**Figure 6.**
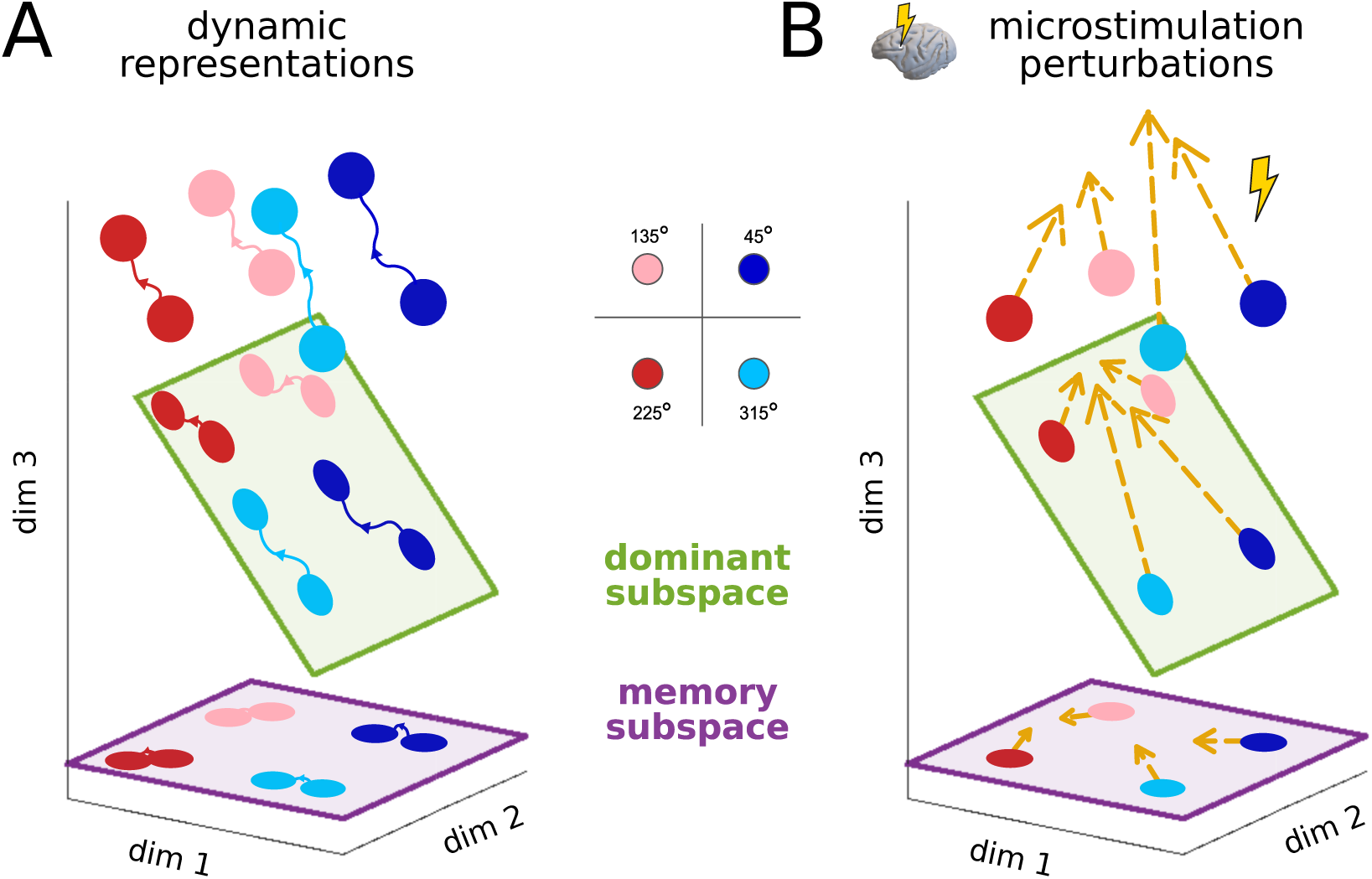
Dissociated population-level mnemonic representations in dlPFC for robustness to perturbations. **A**, Organization of the dominant and memory subspaces in dlPFC’s population activity space. The “dominant” subspace (green) captures the natural evolution of working memory related representations, whereas the “memory” subspace (purple) reflects the stable component of this naturally evolving representation. **B**, Microstimulation perturbations push activity along different directions in neural population activity space. The perturbations are strongly reflected in the dominant subspace (i.e., the perturbations have a substantial projection onto this subspace), but they are minimally reflected in the memory subspace (i.e., the perturbations are largely orthogonal to this subspace).

Several studies that perturbed neural activity in dlPFC during the delay period of working memory tasks were able to induce stronger effects on behavior than the effects reported here. However, these studies involved longer-lasting activity manipulations than the type we performed, e.g., microstimulation throughout the whole delay period (Opris et al., 2005a) or large-scale pharmacological inactivations (Sawaguchi and Iba, 2001). Another related study that showed effects on behavior during working memory applied microstimulation in the frontal eye fields, but also throughout the whole delay period (Opris et al., 2005b). In visually guided tasks, brief microstimulation perturbations during saccade planning have been shown to impact eye movements when applied to the lateral PFC (Wegener et al., 2008) and the dorsomedial PFC (Yang et al., 2008).

In contrast, recent studies that performed transient inactivations of PFC and motor areas during delayed response tasks minimally impacted behavior. In a working memory task, brief optogenetic inactivation of the lateral PFC in monkeys did not impact working memory related signals and behavior (Mendoza-Halliday et al., 2023). In a delayed motor task, transient optogenetic inactivation of mouse premotor cortex (ALM) did not impact behavior (Li et al., 2016; Inagaki et al., 2019). Similarly, in a delayed motor task, transient optogenetic stimulation of monkey motor and premotor cortices produced no behavioral impact, though electrical microstimulation minimally influenced behavior (O’Shea et al., 2022). These studies found that task-relevant signals during the delay period were not affected (Mendoza-Halliday et al., 2023) or quickly recovered (Li et al., 2016; Inagaki et al., 2019) from such transient manipulations, even though activity was strongly modulated by the stimulation. Similarly, we found task-relevant signals that were either not affected by the microstimulation or that tended to recover from it, although they did not always do so (Fig. 5).

Taken together, this body of work points to some possible explanations for how robustness to perturbations is maintained in neural circuits. One possibility is that task-relevant information is affected by the stimulation, but that compensatory mechanisms are in place that restore this information, potentially through redundancy (for example, across hemispheres as in Li et al., 2016). A second possibility is that task-relevant information is affected, but not completely disrupted, and that the information is still accessible in certain subspaces of the same neural population (Murray et al., 2017; Parthasarathy et al., 2019). A third possibility, perhaps related to the second, is that task-relevant information is affected, but that the dimensions where this information is found do not align with the modes of the activity that are truly consequential for behavior (O’Shea et al., 2022). Future work involving additional perturbations and monitoring of neural activity more broadly across cortical areas will be necessary to resolve these potential explanations.

One explicit mechanism by which the brain may retain robustness in the face of perturbation is an attractor network. Attractors are robust to brief and modest perturbations, such as noise, sensory distractors, or direct activity manipulations (Wang, 2021). There are multiple attractor structures that might underlie the maintenance of working memory related information in dlPFC and that could implement the possibilities discussed above (Li et al., 2016; Inagaki et al., 2019; Parthasarathy et al., 2019; Wang, 2021; Zhou et al., 2023; Stroud et al., 2024). Our results are not consistent with multistability (multiple discrete attractors), given that perturbations strongly pushed activity toward different target angle locations but activity did not settle in those locations (Fig. 5A). Furthermore, the monkeys did not tend to saccade toward an incorrect target location. In contrast, evidence for multistability was found in the mouse ALM during delayed response tasks using optogenetic perturbations (Inagaki et al., 2019). We also observed that the perturbed trajectories tended to return to the end point of the unperturbed trajectory (Fig. 5A), and not to an intermediate position along the unperturbed trajectory, which is consistent with point attractors. In contrast, in mouse ALM during delayed motor tasks (Li et al., 2016) and in artificial RNNs trained in memory tasks (Zhou et al., 2023), perturbed trajectories always converged back to the unperturbed trajectory, and not the end point. This might be consistent with input driven dynamics restoring information from interconnected areas such as thalamus (Sauerbrei et al., 2020) or redundant modules across hemispheres (Li et al., 2016), or might indicate the presence of dynamic attractors (Laje and Buonomano, 2013; Zhou et al., 2023).

The work presented here aimed to understand the response of PFC to microstimulation perturbations, and found that working memory signals were robust to such perturbations. An important goal for future work is to design perturbations that can selectively impact working memory. One possible approach is to design stimulation experiments to specifically impact targeted neurons and behavior. For example, microstimulation in FEF improves stimulus detection at a location that matches the response field of the neuron that is microstimulated (Moore and Fallah, 2004). Another approach is to optimize stimulation patterns to produce customized effects on neural populations and behavior. This has been attempted with model-driven and closed-loop optimization of stimulation parameters, which is necessary to deal with the large potential search space (Tafazoli et al., 2020; Nejatbakhsh et al., 2023; Minai et al., 2024). Understanding the ways in which populations of neurons maintain robust cognitive states, along with the development of tools to customize perturbations of neural populations to achieve specific targeted states, is an important goal of modern systems neuroscience.

## Author contributions

J.S.M., Y.M., M.A.S. and B.M.Y. designed research; J.S.M. and Y.M. performed experiments; J.S.M. analyzed data; All authors actively participated in the interpretation of the data; J.S.M. wrote the original draft; J.S.M., Y.M., M.A.S. and B.M.Y. edited the draft and wrote the final paper.

## Acknowledgements

We are grateful to the members of the Yu, Smith, Chase and Batista labs for valuable discussions, to Samantha Schmitt for assistance with data collection, and to Karen McCracken and Mary Ellen Smyth for animal care.

## Data Availability Statement

The data will be made available upon publication.

## Code Availability Statement

The code to reproduce the analysis will be made available upon publication.

